# *Ralstonia solanacearum* depends on catabolism of myo-inositol, sucrose, and trehalose for virulence in an infection stage-dependent manner

**DOI:** 10.1101/700351

**Authors:** Corri D. Hamilton, Olivia Steidl, April M. MacIntyre, Caitilyn Allen

**Affiliations:** Department of Plant Pathology University of Wisconsin-Madison, 1630 Linden Drive Madison, Wisconsin 53706 USA; Microbiology Doctoral Training Program, University of Wisconsin-Madison

**Keywords:** bacterial wilt disease, xylem sap, bacterial catabolism, sucrose, trehalose, myo-inositol, metabolic trade-offs

## Abstract

The soilborne pathogen *Ralstonia solanacearum (Rs)* causes lethal bacterial wilt disease of tomato and many other crops by infecting host roots and then colonizing the xylem vessels. Tomato xylem sap is nutritionally limiting but it does contain sucrose and trehalose. Transcriptomic analyses revealed that *Rs* expresses distinct sets of catabolic pathways at low cell density (LCD) and high cell density (HCD). To investigate the links between bacterial catabolism, infection stage, and virulence, we measured the *in planta* fitness of bacterial mutants lacking carbon catabolic pathways expressed at either LCD or HCD. We hypothesized that the bacterium needs LCD carbon sources early in disease (root infection) while HCD carbon sources are required during late disease (stem colonization). An *Rs ΔiolG* mutant unable to use the LCD nutrient myo-inositol was defective in root colonization but once it reached the stem, this strain colonized and caused symptoms as well as wild type. In contrast, *Rs* mutants unable to use sucrose (*ΔscrA)*, trehalose (*ΔtreA)*, or both *(ΔscrA/treA*), infected roots as well as wild type but were defective in colonization and competitive fitness in tomato mid-stems and were reduced in bacterial wilt virulence. Additionally, xylem sap from tomato plants colonized by *ΔscrA, ΔtreA, or ΔscrA/treA* contained more sucrose than sap from plants colonized by wild-type *Rs*. Together, these findings suggest *Rs* metabolism is specifically adapted for success in the different nutritional environments of plant roots and xylem sap.

---

Microbial success hinges on the ability to be niche adapted and reduce organic compounds. When facing niches with diverse and scares nutrient availabilities having a large scope of catabolism pathways is essential (Fatima & Senthil-Kumar, 2015, Siebrecht et. al., 2003). However, there is a cellular cost for utilizing large numbers of catabolic pathways (Dekel & Alon, 2005, Polz & Cordero et. al. 2018, Pfeiffer & Bonhoeffer, 2004, Gudelj et. al., 2007). Microbes use many signaling mechanisms to make informed choices about what pathways to activate and when (Wadhams & Armitage, 2004). There is a careful balance between wasting energy from improper pathway regulation and gaining energy from the consumption of carbon sources. Growth and reproduction of all living organisms relies on the optimization of these chemical interactions with their environment.

Myo-inositol, sucrose, and trehalose are all compounds that influence the plant host environment for many plant-associated microbes. Myo-inositol is a carbon source often found in high quantities *in plant*a and is the most abundant carbon source found in root nodules formed by association of Bradyrhizobium diazoefficiens and host pea (Sköt and Egsgaard 1984). The first of many nutrient interfaces for plant-associated microbes is root exudates where myo-Inositol is commonly found (Wang & Bergeson, 1974, Wei et. al., 2015). However, sucrose is the major form of plant carbon produced through photosynthesis (Salanoubat & Belliard, 1989). It is continuously shuttled from source to sink and is essential for plant growth and development (Pollock & Farrar, 1996, Sonnewald & Willmitzer, 1992). Pathogens can modulate the host metabolism and cause plants to produce and reallocate carbon sources like sucrose (Thines et al., 2000). Plants use trehalose signaling to balance nutrient levels, for example trehalose 6-phoshate influences *Arabidopsis’* sucrose and starch ratios in left carbon storage systems (Kolbe et al. 2005). Trehalose is well characterize for its roles in plant metabolic regulations and stress tolerance (Lunn et al., 2014, Paul et. al., 2008). Myo-inositol, sucrose, and trehalose are all compounds the many microbes can consume and could modulate host plant environment conditions to be more favorable for invading pathogens.

*Ralstonia solanacearum (Rs)* is part of a heterogeneous species complex of plant pathogenic β-proteobacteria that cause bacterial wilt disease (BWD) (Allen et. al., 2005, Genin & Denny, 2012, Prior et. al., 2016) This pathogen indiscriminately stifles yield for subsistence and commercial crops (Elphinstone, 2005). The pathogen’s wide host range and high environmental survival contribute to the large losses seen each year. *Rs* causes disease on over 100 economically relevant crops including tomato, potato, banana, tobacco, peanut, ginger, cloves, and eggplant (Allen et. al., 2005). To investigate the *Rs*-tomato pathosystem we used the Asian strain GMI1000, belonging to phylotype I sequevar 18 and causes major losses to tomato growers (Prior et. al., 2016).

*Rs* spends its life cycling between two dynamic nutritional environments: rhizosphere soil and xylem sap. The rhizosphere is teeming with a wide variety of microbes and carbon sources (Yang & Crowley, 2000). *Rs* is well adapted to surviving in this environment with >50 catabolic pathways to scavenge the many available resources, although specific pathway profiles vary within the species complex (Álvarez, B. et al., 2010, Jacobs et. al., 2012, Zuluaga et. al. 2013). *Rs* remains a relatively low titers in the bulk soil of infected fields around <10^4^ CFU/gram (Allen et. al., 2005, Hayward, 1991). *Rs* senses and chemotaxes to locate and enter plant roots via wounds or natural openings.

We know that *Rs* metabolic pathways are differentially regulated during infection with *Rs ΔphcA*, a quorum sensing mutant that is a proxy for the low cell density (LCD) early infection regulatory mode. PhcA is a global regulator that mediates a trade-off from maximizing growth to producing costly virulence factors (Khokhani et. al., 2017, Polz & Cordero et. al. 2018) Interestingly, these results suggest that *Rs* uses a set of carbon sources available in early infection (e.g. myo-inositol) but shifts preferences during late infection or high cell density (HCD) (e.g. sucrose)(Jacobs et. al. 2012, Khokhani et. al., 2017, Lowe-Power et. al., 2018). Looking at these dynamic shifts in *Rs* behavior in plant xylem sap narrowed the search for carbon sources contributing to *Rs* virulence and colonization in specific infection stages.

Within 24 hours *Rs* will travel through the developing vascular bundles to reach the xylem vessels of susceptible tomato roots (McGarvey et. al., 1999, Caldwell et. al. 2017). Once inside a host, *Rs* occupies the xylem sap, historically described as a nutrient-poor solution of water and minerals. The vascular vessels themselves are dead, lacking the potential nutrient sources found in living cells (Evert & Eichhorn, 2006). The xylem vascular system acts as a highway for *Rs* to spread systemically through its host. *Rs* promptly multiplies to >10^9^ CFU/gram stem and eventually grows into aggregates in a biofilm matrix that can fill entire vessels and obstruct water flow (Gnanamanickam, 2006) The completion of the *Rs* life cycle results in the common symptoms of BWD, including wilting of the leaves, petiole epinasty, vascular discoloring, rot, and death.

Little is understood about how *Rs* grows to such high cell densities in a relatively harsh, nutrient-poor environment like xylem sap. We now know that tomato xylem sap contains enough sugars, amino acids, and organic acids to support limited bacterial growth *in vitro* (Coplin et. al., 1974, Dixon & Pegg et. al., 1972, Evert & Eichhorn, 2006, Fatima & Senthil-Kumar, 2015, Wang & Bergeson, 1974, White et. al., 1981). These nutrients vary by plant species, season, time of day, and growing conditions (Siebrecht et. al., 2003, Zuluaga et. al., 2013). Infected xylem sap is enriched in metabolites, including many carbon sources that support *Rs* growth (Lowe-Power et. al., 2018). This metabolomics finding suggests that the presence of *Rs* increases accessible xylem carbon. A transcriptomic analysis shows that several metabolic pathways in *Rs* are differentially regulated during infection (Jacobs et. al., 2012). This work advocates that sucrose, trehalose could fuel *Rs* growth *in planta*.

Our objective was to elucidate the links between *Rs* catabolism, infection stage, and ability to colonize and cause disease on tomato crops. We hypothesize that *Rs* is adapted to two dynamic nutritional environments: rhizosphere soil and xylem sap by using specific carbon sources at different stages in infection cycle. We analysis bacterial fitness compare to wildtype using bacteria catabolic mutants knocking out *Rs* ability to consume myo-inositol, sucrose, trehalose, and sucrose & trehalose (*ΔiolG, ΔscrA, ΔtreA, and ΔscrA/treA)*. We focused on identifying if specific carbon sources that *Rs* consumes required during BWD. We found that LCD gene (iolG) is required for early/root infection and HCD genes (scrA & treA) are required for late/stem infection. When determine through enzymatic assays that sucrose is require for consumption whereas trehalose is modulating plant response rather that being a primary carbon source for bacterial growth in late infection.

## RESULTS

### ‘Omics analysis and phenotypic characterization of bacterial catabolic genes, iolG, scrA, and treA

*Rs (GMI1000)* bacterial genes iolG, scrA, and treA are functional *in planta*, absolute expression being 10.15, 11.08, and 12.53 respectively (Jacobs et. al., 2012). Bacterial gene *iolG* encodes an oxidoreductase 2-dehydrogenase signal peptide protein in the myo-inositol catabolism pathway. *scrA* encodes a sucrose-specific transmembrane protein or sucrose importer. *treA* encodes an alpha-trehalase signal peptide protein associated with the trehalose catabolism genes. Myo-inositol, sucrose, and trehalose can be used by GMI1000 as a sole carbon source, detected in *ex vivo* xylem sap, and found in root exudates (Lowe-power et. al.2016, Wang & Bergeson, 1974, Wei et. al., 2015). Tomato cultivar Bonny Best *ex vivo* xylem sap metabolites sucrose and myo-inositol are increased less than 2-fold change whereas trehalose is increase 19-fold change between healthy and GMI1000 Infected sap (Lowe-Power et. al. 2016). An *in planta* transcriptomic analysis between *ΔphcA*, a quorum sensing mutant that is a proxy for a LCD locked state. *IolG* is up regulated 2.08-fold in the phcA background compared to wildtype corresponding with myo-inositol consumption being up regulated at LCD. treA and scrA in the phcA background are both down regulated 2.92 and .54-fold compare to wildtype corresponding with trehalose and sucrose consumption being down regulated at LCD (Khokhani et. al., 2017) (Table 1). To determine whether *ΔiolG, ΔscrA, and ΔtreA* were phenotypic knockouts of our target catabolism pathways. Each strain and complement was made through double homologous recombination to create in-frame deletions or insertions using primers listed in Supplemental Table 1. To determine the metabolic profile of each mutant and complement, their growth fitness was recorded when grown on sole carbon sources of glucose, myo-inositol, sucrose, and trehalose. *ΔiolG, ΔscrA, and ΔtreA, ΔscrA/treA* all grow like wildtype on glucose. *ΔiolG, ΔscrA, ΔtreA and ΔscrA/treA* differed from wildtype growth only in their respective pathways myo-inositol, sucrose, trehalose, or both sucrose and trehalose. Additionally, all complements restored mutant growth phenotype to wildtype or better than mutant (Table 2). This confirmed that in fact the mutants used in this investigation were in fact functional and specific catabolic gene deletions.

**Table 1:** *R. solanacearum* catabolic genes of interest.

| Carbon Source | Gene Name | <sup>1</sup>In planta<br>LogFC<br>WT vs. *ΔphcA* | <sup>2</sup>In plant<br>Absolute<br>Expression | <sup>3</sup>Metabolite<br>Fold Change<br>(Infected/Healthy) | <sup>4,5</sup>Gene Annotation | Locus |
| --- | --- | --- | --- | --- | --- | --- |
| Myo-inositol | *iolG* | 2.08 | 10.15 | 1.95 | Oxidoreductase 2-dehydrogenase signal peptide protein | Rsc1246 |
| Sucrose | *scrA* | -0.54 | 11.08 | 1.29 | Sucrose-specific transmembrane protein | RSc1285 |
| Trehalose | *treA* | -2.92 | 12.53 | 19.39 | Alpha-trehalase signal peptide protein | RSp 0277 |

<sup>1</sup> FC indicates gene expression fold change values in *ΔphcA* relative to wild-type as described in Khokhani *et. al.* 2017.
<sup>2</sup> Gene absolute expression values were based on four biological replicates, normalized using the Robust Multichip Algorithm as described in Jacobs *et. al.* 2012.
<sup>3</sup> Values represent the Bonny Best *ex vivo* xylem sap metabolite fold change between Healthy and *Ralstonia* (GMI1000) Infected sap as described in Lowe-Power *et. al.* 2016.
<sup>4</sup> Gene annotations based on www.ncbi.nlm.nih.gov/nuccore/NC_003295.1 and www.ncbi.nlm.nih.gov/nuccore/NC_003296.1.
<sup>5</sup> Alignment of carbon sources, locus, and pathways based on KEGG.

**Table 2:**
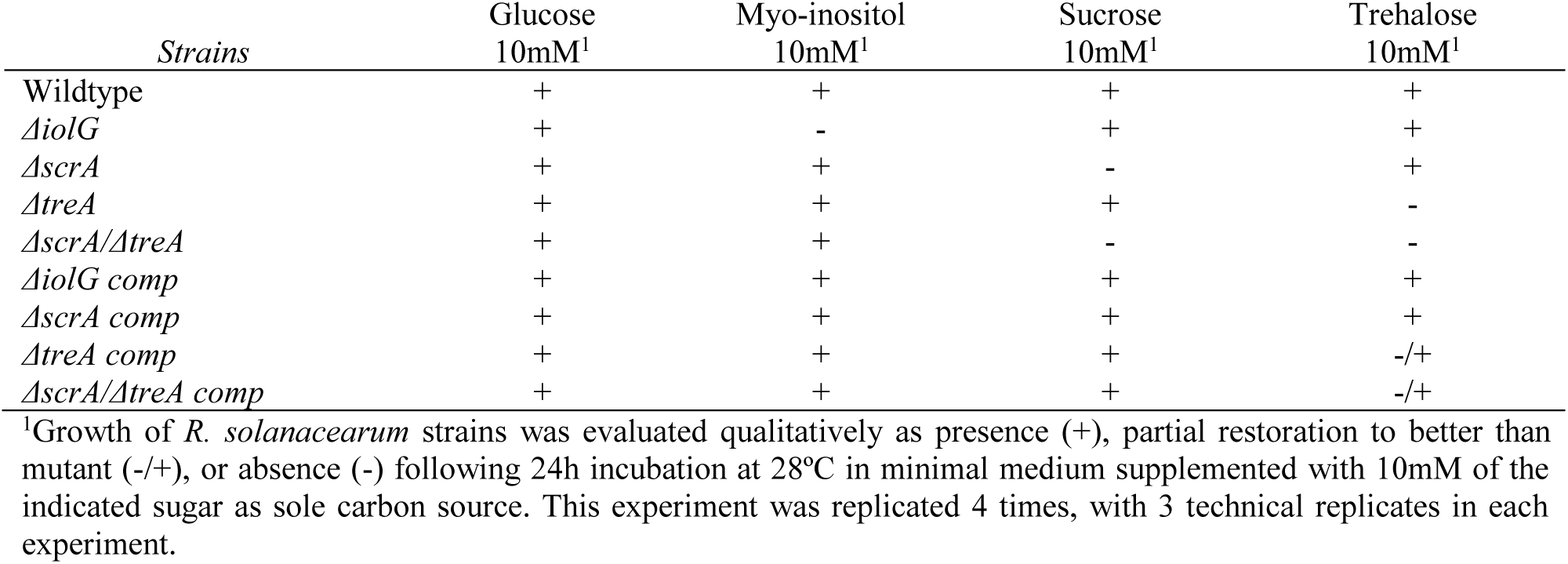
Growth of *R. solanacearum* catabolic mutants on relevant sole carbon sources.

| Strains | Glucose<br>10mM<sup>1</sup> | Myo-inositol<br>10mM<sup>1</sup> | Sucrose<br>10mM<sup>1</sup> | Trehalose<br>10mM<sup>1</sup> |
| --- | --- | --- | --- | --- |
| Wildtype | + | + | + | + |
| $\Delta iolG$ | + | - | + | + |
| $\Delta scrA$ | + | + | - | + |
| $\Delta treA$ | + | + | + | - |
| $\Delta scrA/\Delta treA$ | + | + | - | - |
| $\Delta iolG$ comp | + | + | + | + |
| $\Delta scrA$ comp | + | + | + | + |
| $\Delta treA$ comp | + | + | + | -/+ |
| $\Delta scrA/\Delta treA$ comp | + | + | + | -/+ |

<sup>1</sup>Growth of *R. solanacearum* strains was evaluated qualitatively as presence (+), partial restoration to better than mutant (-/+), or absence (-) following 24h incubation at 28°C in minimal medium supplemented with 10mM of the indicated sugar as sole carbon source. This experiment was replicated 4 times, with 3 technical replicates in each experiment.

### Myo-inositol catabolism is required for colonization & competitive fitness in tomato roots

When *R. solanacearum* GMI1000 strains were separately introduced directly into tomato root infection myo-inositol consumption was required for full root infection. Tomato plants were inoculated with wild-type GMI1000, *ΔphcA, ΔscrA, ΔtreA*, or *ΔiolG* onto the roots and the population size of each strain was quantified in each seedling 24 & 72 hours post-inoculation (Fig. 1). The LCD locked mutant *ΔphcA* control grew as expected, doing well early in disease and failing at late disease. We found that genes *scrA, treA, and scrA/treA* are not required during the 1^st^ 24 hours of root colonization. However, at 24 hours the *iolG* mutant is reduced in root colonization compared to wildtype (Mann-Whitney). Within 24 hours *Rs* will travel through the developing vascular bundles to reach the xylem vessels of susceptible tomato roots. This means 72 hours post infection represents stem or late disease (Caldwell, D. et al. 2017, McGarvey JA. et al. 1999). As expected only genes *scrA, treA, and scrA/treA* are required for 72 hours root colonization. This suggest that bacterial myo-inositol consumption is required for early colonization.

**Figure 1:**
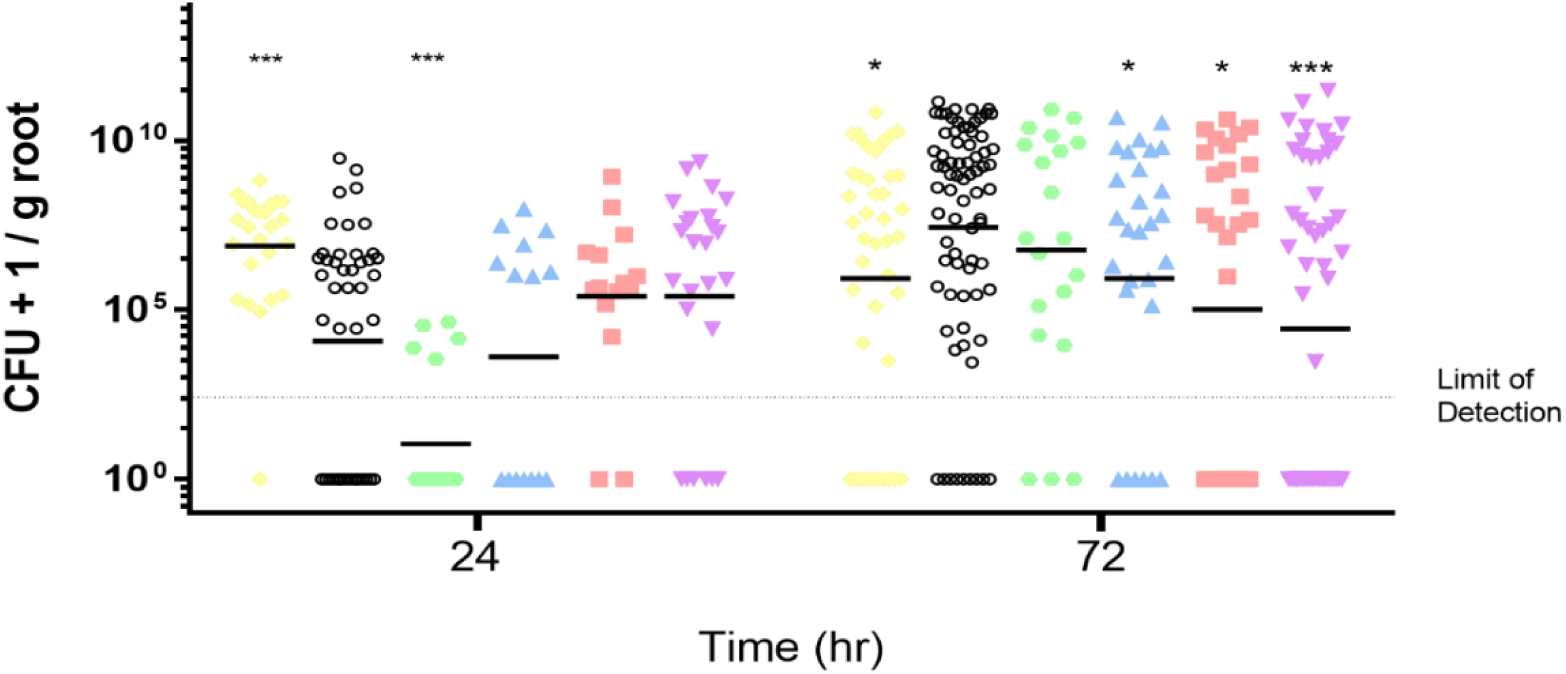
The catabolic mutants they can’t consume myo-inositol (*ΔiolG)* is defective at 24hr while *scrA and treA* mutants are defective at only 72hr. To test the hypothesis that myo-inositol unlike sucrose and trehalose is important for early disease progress. I infected tomato plants with a GmR-marked *ΔphcA*, KmR-marked wild-type(black), markerless *ΔiolG(green)*, markerless *ΔscrA(blue)*, SpecR-marked *ΔtreA(red)*, and SpecR-marked *ΔscrA/treA(purple)* bacteria by root attachment inoculation of 4-day old BB tomato seedlings. *Rs* population sizes were quantified by serial dilution plating of ground 2 root sections harvested 24, 48, 72 hours after inoculation. This confirms the hypothesis that the *ΔscrA, ΔtreA, & ΔscrA/treA* mutants can colonize tomato seedling as well as wild-type and *ΔiolG* mutant final to colonize tomato seedling as well as wild-type. *ΔphcA* is used as a low cell density lock mutant control expected to do well at early infection only. Data shown represent 3-6 biological replications, each containing 12-20 plants per treatment, and Horizonal black line represents geometric mean of each strain. Significance by Mann-Whitney ^a^ p<.001, ^b^ p<.0001, & ^c^ p<.00001

To specifically measure the contribution of bacterial catabolism to early infection success, we used an *in planta* root competition assay to determine whether the GMI1000 bacterial catabolic mutants was reduced in competitive fitness. Tomato plants were inoculated with a 1:1 mixture of wild-type GMI1000 and GMI1000 *ΔscrA, ΔtreA, ΔiolG* directly onto roots and the population size of each strain was quantified in each plant 24 & 72 hours post-inoculation (tab. 3). Only the *ΔiolG*, mutant has reduced colonization compared to wildtype (Mann-Whitney). This indicating that only the ability to catabolize root available myo-inositol provides *Rs* with a significant fitness advantage during root infection.

**Table 3:**
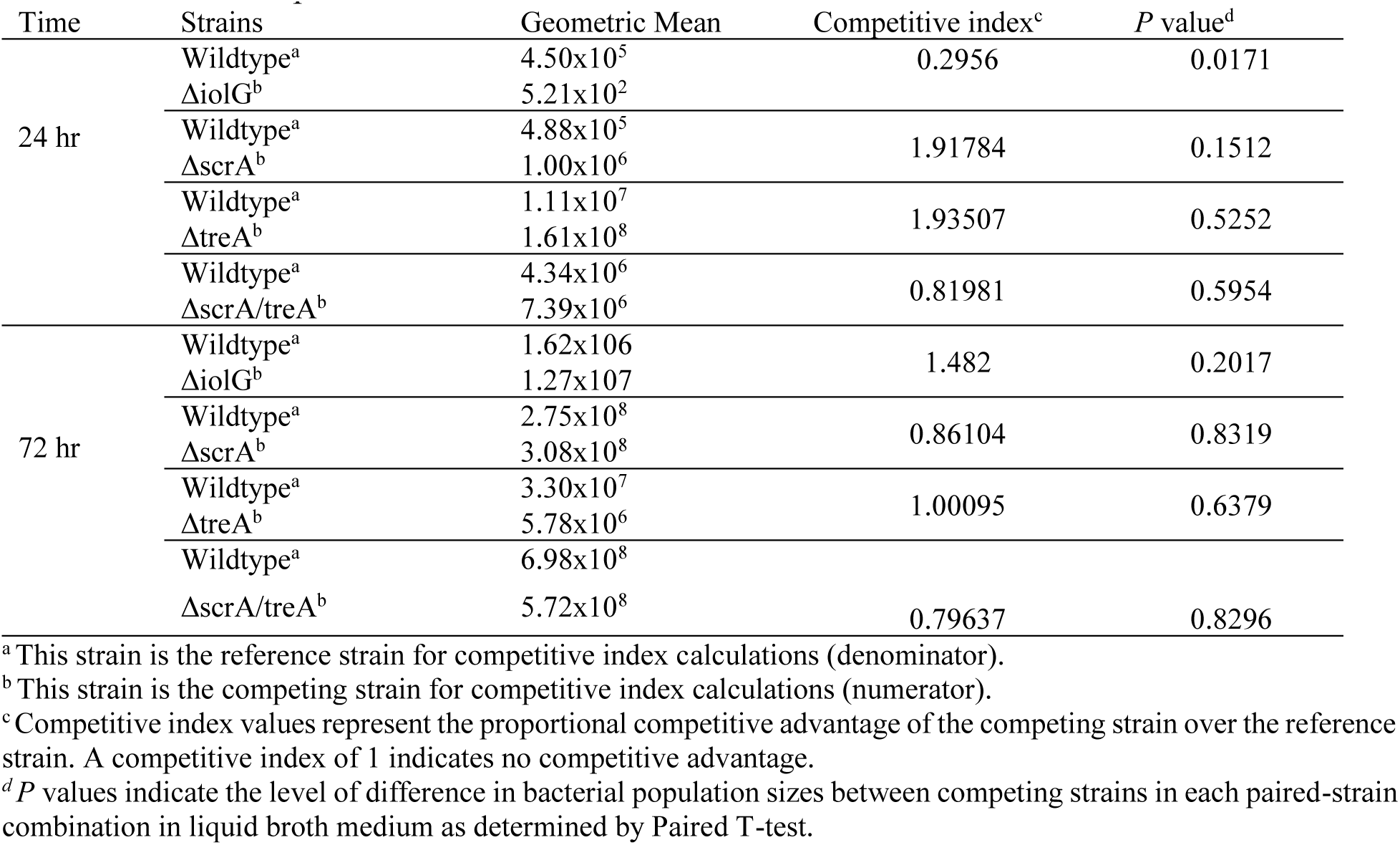
Root Competitive Index.

### *R. solanacearum* requires a scrA, treA, and scrA/treA for full virulence

*Rs* infection modulates xylem sap conditions during infection to increase *Rs* available carbon sources like sucrose. We determined strain effects of virulence by individually inoculating each strain of bacteria on to susceptible cultivar Bonny Best tomato plant and rated disease index over 14 day (Fig 2.). We found that *ΔiolG* is equally as virulent as wildtype GMI1000.

**Figure 2:**
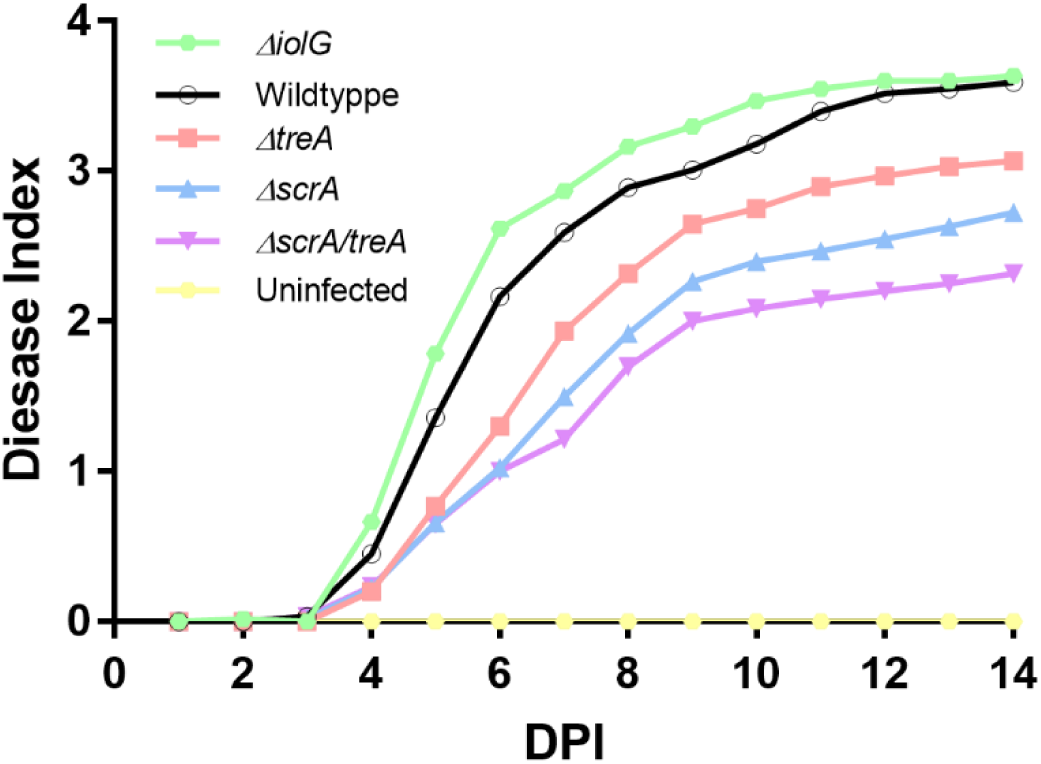
*R. solanacearum* needs catabolism of sucrose and trehalose for full bacterial wilt virulence. Unwounded 17-day-old tomato plants (wilt-susceptible cv. Bonny Best) were inoculated by pouring a dilute bacterial suspension into the pot to a final pathogen concentration of 5×10^7^ CFU/gram potting mix. Plants were incubated in a 28°C growth chamber and symptoms were rated daily on a 0-4 disease index. Data shown are the mean of four (wild-type n=8) independent experiments, each containing 15 plants per treatment. Mutants *ΔtreA, ΔscrA*, and *ΔscrA/treA* all have lower virulence than the wild-type (P< 0.002, repeated measures ANOVA). Virulence of the *ΔiolG* catabolic mutant is not different from wild-type (P> 0.05, repeated measures ANOVA).

However, single mutants *ΔscrA & ΔtreA* are both defective in virulence compared to wildtype. The double mutant *ΔscrA/treA* virulence is reduced more than both single mutant parents. This suggest that *ΔscrA, ΔtreA, & ΔscrA/treA* are required for full virulence and that there is some additives virulence defect with the double mutant.

### Sucrose catabolism is required for colonization & competitive fitness in tomato stems

When *R. solanacearum* GMI1000 strains were separately introduced directly into tomato vascular systems via a soil soak. Tomato plants were inoculated with wild-type GMI1000, *ΔscrA, ΔtreA*, or *ΔiolG* into the xylem through soil drench inoculation and the population size of each strain was quantified in each plant 5 days post-inoculation (Fig. 3). We found that genes *iolG* and *treA* are not required for stem colonization. The *ΔscrA* and *ΔscrA/treA* mutants have reduced colonization compared to wildtype (Mann-Whitney). Again, the double mutant *ΔscrA/treA* colonization is reduced more than both single mutant parents. This suggest that *ΔscrA & ΔscrA/treA* are required for stem colonization and that there is some additives virulence defect with the double mutant.

**Figure 3:**
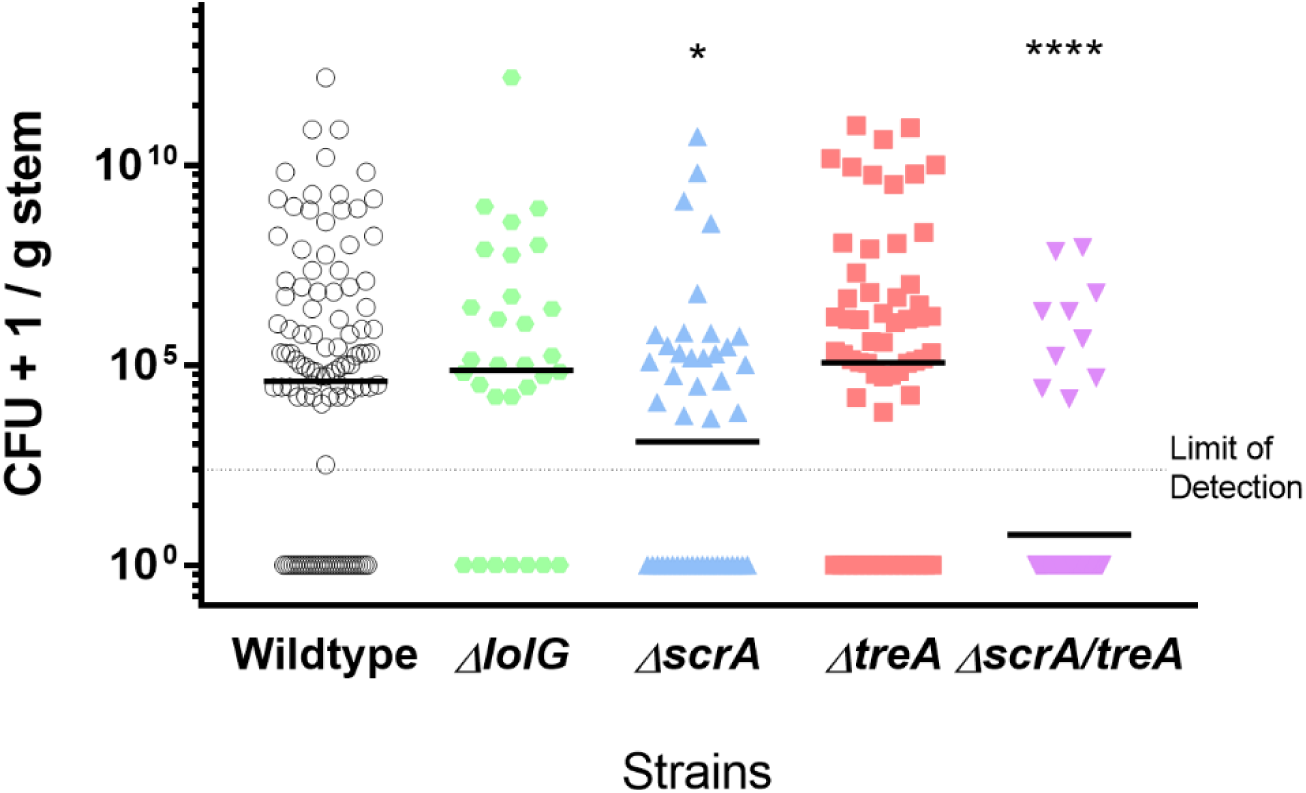
Sucrose catabolism helps *R. solanacearum* colonize tomato stems. Unwounded 17-day-old tomato plants wilt-susceptible cv. Bonny Best were inoculated by soil-soaking with 5×10^7^ CFU/gram bacterial suspension of KmR-marked wild-type(black), markerless *ΔiolG(purple)*, markerless *ΔscrA(blue)*, SpecR-marked *ΔtreA(red)*, and SpecR-marked *ΔscrA/treA(green)* bacteria. After 5 days the bacterial population size in each plant was determined by dilution plating a ground 0.1 g stem section harvested just above the cotyledon. Data shown from 4 independent experiments, each containing 12-20 plants per treatment. Horizonal black line represents geometric mean of each strain. Asterisk indicates differences from wild-type (P< 0.05, one asterisk; P<0.00001, 4 asterisks, Mann-Whitney)

To determine the contribution of xylem carbon sources to bacterial stem success, we used an *in planta* stem competition assay to determine whether the GMI1000 bacterial catabolic mutants were reduced in competitive fitness. Tomato plants were inoculated with a 1:1 mixture of 2000 cell of wild-type GMI1000 and GMI1000 *ΔscrA, ΔtreA, ΔiolG* directly into the xylem through cut petioles, and the population size of each strain was quantified in each plant 120 hours post-inoculation (Tab. 4). The *ΔscrA, ΔtreA* and *ΔscrA/treA* mutants have reduces colonization compared to wildtype (Mann-Whitney). This indicating that the ability to metabolize xylem sucrose provides *R. solanacearum* with a significant fitness advantage during stem infection.

**Table 4:** Competitive fitness of *R. solanacearum* catabolic mutants in tomato midstems, relative to wild type.

| Time (hr) | Strains | CFU/g stem<br>(Geometric Mean) | Competitive<br>index<sup>c</sup> | *P* value<sup>d</sup> |
| --- | --- | --- | --- | --- |
| 120 | Wildtype<sup>ae</sup> | 3.28E+07 | 0.9483 | 0.3274 |
| | ΔiolG<sup>be</sup> | 4.79E+07 | | |
|  | Wildtype<sup>a</sup> | 4.10E+09 | 0.5676 | 0.0002 |
| | ΔscrA<sup>b</sup> | 7.65E+09 | | |
|  | Wildtype<sup>a</sup> | 1.09E+09 | 0.4231 | 0.0009 |
| | ΔtreA<sup>b</sup> | 4.50E+08 | | |
|  | Wildtype<sup>a</sup> | 3.47E+05 | 0.5000 | 0.0123 |
| | ΔscrA/treA<sup>b</sup> | 2.78E+05 | | |

<sup>a</sup>This strain is the reference strain for competitive index calculations (denominator).
<sup>b</sup>This strain is the competing strain for competitive index calculations (numerator).
<sup>c</sup>Competitive index values represent the proportional competitive advantage of the competing strain over the reference strain. A competitive index of 1 indicates no competitive advantage.
<sup>d</sup>*P* values indicate the level of difference in bacterial population sizes between competing strains in each paired-strain combination in liquid broth medium as determined by Paired T-test.
<sup>e</sup>Data shown is representative of 3 rather than 4 biological replications

### Bacterial growth based on elevated sucrose concentration in *ex vivo* xylem sap

Excess carbon sources are associated with increased *Rs* growth (Wang & Bergeson, 1974, Fatima & Senthil-Kumar, 2015, Siebrecht et. al., 2003). GMI1000 infected *ex vivo* xylem sap has increase metabolites compared to healthy ex vivo xylem sap (Lowe-Power et. al., 2016). To determine if catabolic mutants leave more metabolites for *Rs* consumption compared to wildtype, we use previously wildtype or mutant infected plant xylem sap as growth media for wildtype, *ΔiolG, ΔscrA, ΔtreA, & ΔscrA/treA* (Fig. 4). All strains grow the same on sap previously infected with *ΔiolG and ΔtreA compared to* sap previously infected with wildtype. Most strains grow better on previously infected with *ΔscrA and ΔscrA/treA sap* except the *ΔscrA* & *ΔscrA/treA* mutant. This suggest that at least sucrose is modulating bacterial growth in ex vivo xylem sap. Interesting *ΔscrA/treA sap* seem to support more growth than *ΔscrA* or *ΔtreA* alone.

**Figure 4:**
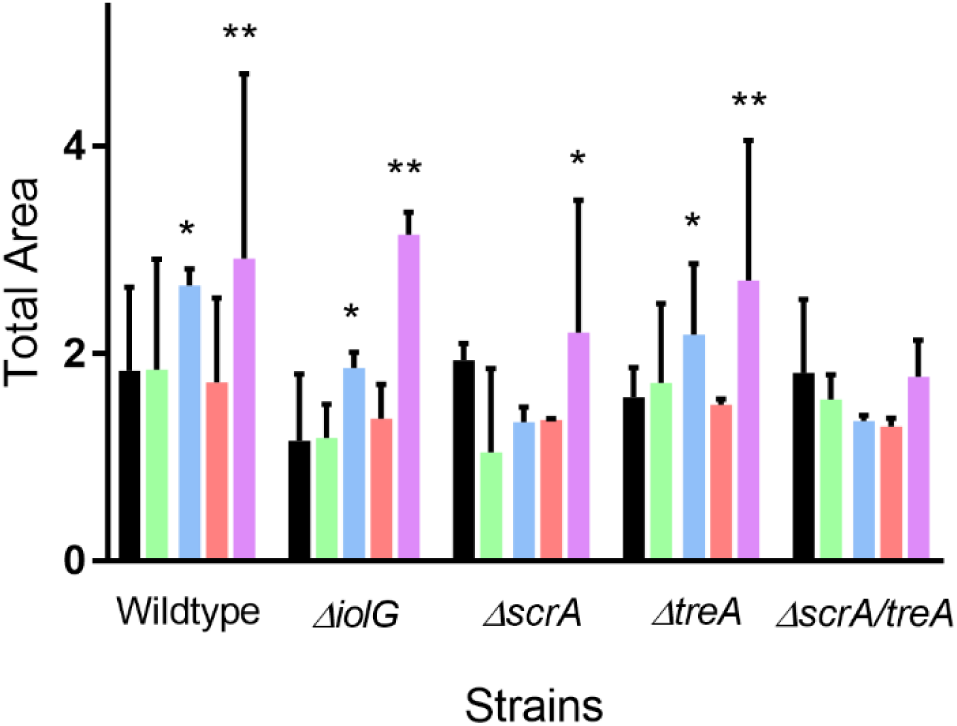
*Ex vivo* xylem sap from plants previously infected with ScrA, ScrA/TreA mutants increase ex vivo xylem sap growth of *R. solanacearum* strains that can consume sucrose. Growth of R. solanacearum strains was evaluated quantitively by calculating Area Under the curve as following 36h incubation at 28°C in ex vivo xylem sap. Rs strains were grown on sap pre-infected by wildtype (black), ΔiolG (green), ΔscrA (blue), ΔtreA (red), & ΔscrA/treA (purple). This experiment was replicated 3-6 times, with 3 technical replicates in each experiment. (P< 0.06, one asterisk; P<0.01, 2 asterisks, Mann-Whitney)

To investigate the concentration of sucrose left in *ex vivo* xylem sap we use an enzymatic assay of previously infected xylem sap (Fig. 5). Wildtype and *ΔiolG* both have the same level of sucrose while *ΔscrA & ΔscrA/treA* have increased levels of sucrose compared to previously infected with wildtype sap. *ΔtreA* unexpectantly also increase sucrose levels equal to *ΔscrA/treA.* Although previously infect *ΔscrA/treA sap* induces the highest wildtype growth, *ΔscrA* sap contains the highest sucrose concentration. Suggesting that an increase in sucrose is responsible for some increase growth and that the additive effect of treA and scrA deletions is not fully explained by increase in sucrose concentration.

**Figure 5:**
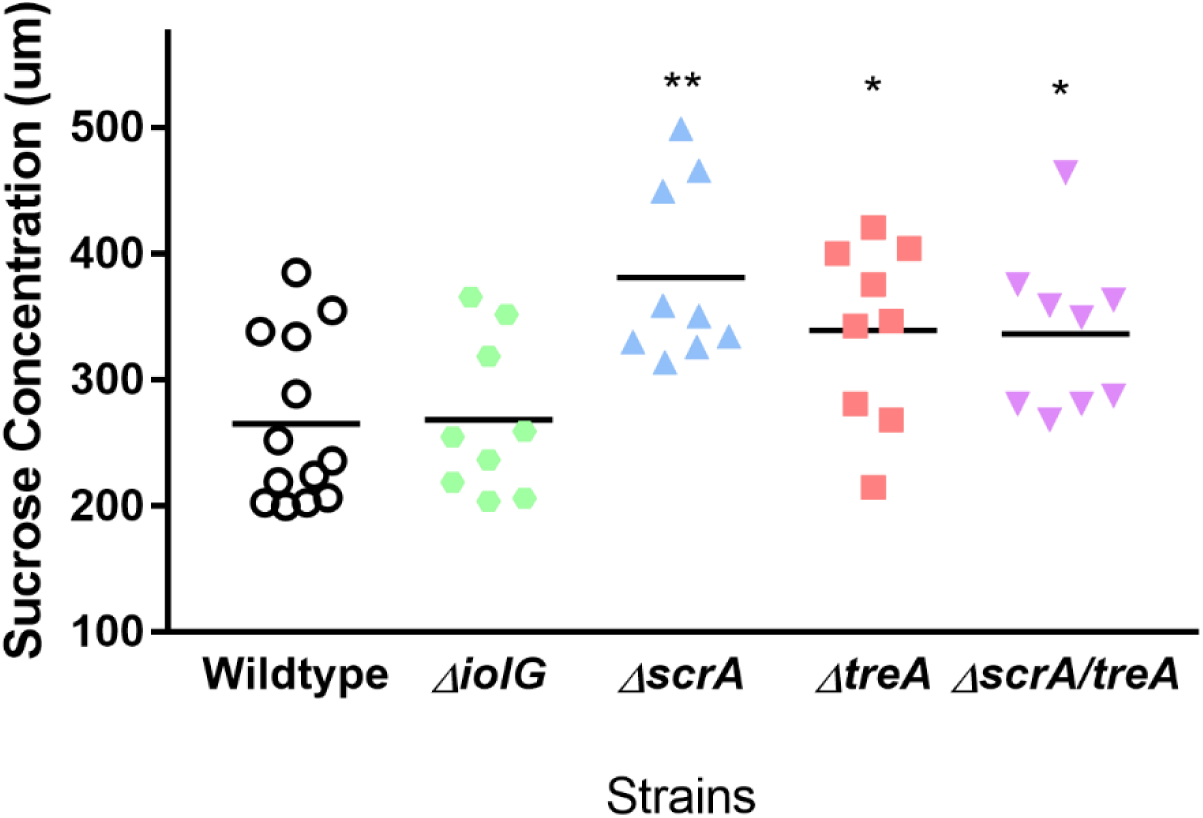
ScrA, TreA, and ScrA/treA mutants modulate sucrose levels in xylem. Sucrose quantification by the GOD/invertase method. Standard sucrose solutions were also added to each plate in separate wells at concentrations 10-1000 umol sucrose. This allowed a calibration curve to be constructed which was used to determine the sucrose concentration in each sample. Each point represents the means of triplicate technical replication and analysis was done in triplicate biological replications. Significance by Student-Paired T-Test, *p<.01, **p<.001.

## Discussion

### Nutrient varied environments during pathogenesis

To understand how *Rs* can survive and infect host in two distinct environments, we needed to understand what carbon sources are required for the pathogen to be successful in each environment. The soil or root environment has been characterized as an environment teeming with small quantities of a diverse set of nutrient sources and high levels of competition (Fatima & Senthil-Kumar, 2015, Siebrecht et. al., 2003, Bais et. al., 2006). Recently we’ve learned about the nutrient availability in the highly selective xylem environment. Xylem sap has *Rs* available carbon sources and that *Rs* infection increases the concentration of several xylem metabolites (Lowe-Power et. al. 2016, Jacobs *et al*., 2012, *Dalsing et al*., 2015, Coplin & Sequeira *et al*., 1974, Chellemi *et al*., 1998, Zuluaga *et al*., 2013.)

We extend our understanding of the role of pathogen utilization of carbon sources in virulence by creating a series of deletion mutants in catabolic pathways, including *iolG, scrA*, and *treA*. These mutants were specifically targeted because they represent LCD or HCD bacterial upregulated pathways, a sole carbon source for *Rs*, and is found in plant root exudates or xylem sap. *Rs* uses a set of carbon sources available in early infection including myo-inositol. Then *Rs* shifts preferences during late infection or high cell density (HCD) to sucrose consumption.

### *Rs* myo-inositol consumption is required for root colonization

We found that catabolism of myo-inositol is required for *Rs* infection and colonization of tomato roots by utilizing information gained from comparing the transcriptome of *ΔphcA* and wildtype GMI1000 during infection. PhcA regulator mediates a tradeoff from a broad metabolic capacity at LCD to a much narrower capacity at HCD (Khokhani et. al., 2017 & Bais et. al., 2006). This shift mediated by PhcA parallels the shift between early and late-stage plant infections.

Unsurprisingly *ΔphcA*, a global regulator mutant, is avirulent (Khokhani et. al., 2017, Lowe-Power et. al., 2018b). However, we found high populations of this mutant in root colonization assays. A *ΔphcB* mutant, which is unable to synthesize the 3-OH-MAME quorum sensing signal and thus quorum insensitive, grew like *ΔphcA* on various sole carbon sources *in vitro* (supplemental fig. 1). This confirm that *ΔphcA* is a good proxy for early and not late infection and that LCD catabolism profiles are conserved across several types of LCD lock mutations.

The myo-inositol pathway upregulated at LCD meaning our bacterial catabolic mutant unable to consume myo-inositol(*ΔiolG*) is important for LCD conditions. We reported that *ΔiolG* is reduced in root colonization but recovers and is not reduced in stem virulence. In tomato root exudate myo-inositol is induced in response to other soilborne pathogens, this could contribute to the plant pathogen defense (Kravchenko et. al., 2003, Cepulyte et. al., 2018). Even though myo-inositol is available in plant xylem sap, bacterial expression increases during early disease progress. This suggest that bacterial myo-inositol consumption is only required for root infection, even though myo-inositol is abundant in xylem sap.

### *Rs s*ucrose consumption is required for full virulence

Sucrose catabolism is not required for *Rs* infection and colonization of tomato roots. We confirmed that LCD gene *iolG* is required for early/root infection and by knocking out bacterial sucrose consumption we determined if HCD genes are required for late/stem infection. We compared ability of *Rs* catabolic mutants to colonize the interior of tomato seedling roots 24 & 72 hpi. Only at the 72-hour time point did lacking sucrose catabolism result in reduced root colonization. By 72 hpi, wilting symptoms were already visible in the tomato seedlings, corresponding to late disease (supplemental fig. 2). This result suggests that sucrose as a food source isn’t necessary for colonization during early stages of disease but plays a role in later disease. Similarly, root competition assays confirmed *ΔscrA* is just as competitive as wildtype during early infection Overall, our data suggest that during early infection at the root, bacterial fitness is not enhanced by catabolism of sucrose.

We identified that *Rs* strain GMI1000 requires sucrose catabolism to fully wilt and kill tomato plants. The single mutants *ΔscrA* had a virulence defect compared to the wildtype strain. Quantification of *ΔscrA* colonization in mid-stem is reduced compared to wildtype. When mutant strains were co-inoculated with their wildtype marked parent directly into tomato plant mid-stem, wildtype GMI1000 outcompeted *ΔscrA*. This provides evidence that sucrose pathway is essential for virulence. To confirm that sucrose catabolism is important our enzymatic assay showed that sucrose is increase in xylem sap when mutants can’t consume the metabolite. This further supports that sucrose is induced during infection and catabolized by GMI1000 to reach full pathogenesis during mid-stem colonization and infection.

### Trehalose catabolism is modulating plant available carbon

Trehalose catabolism is required for *Rs* infection and colonization of tomato stems but not roots. Root competition assays confirmed *ΔtreA* is just as competitive as wildtype during early infection. Like *scrA, treA* only fails at colonizing seedlings equal to wildtype at 72 hours. This confirms the role of trehalose importance during late disease. The single mutants *ΔtreA* has a stem virulence defect compared to the wildtype strain. This provides evidence that trehalose pathway is essential for virulence.

Missing both trehalose and sucrose together doesn’t affect root colonization or competitive fitness compared to wildtype or parent single mutants during early infection. Double mutant *scrA/treA has reduced seedling colonization* at 72 hours the beginning of late disease. The single mutants *ΔtreA* and *ΔscrA* both had slight defects but the double mutant, missing both pathways, had a significant loss in virulence compared to the wildtype strain. This is confirmation of the importance of trehalose and sucrose for full virulence at HCD.

Recent information confirms that trehalose concentration levels in tomato xylem are so low that it’s unlikely that lack of trehalose consumption is affecting population size (MacIntyre et. al. 2019). In culture 1 mM of trehalose is required to induce measurable growth in *Rs* populations and enzymatic assays with the limit of detection of .1mM trehalose couldn’t detect trehalose in wildtype, scrA, treA, or scrA/treA infected xylem <u>sap</u> (supplement fig. 4). Our result that *ΔtreA* is reduce in mid-stem competitive fitness and not stem colonization aligns with the suggestion that absence of treA is not contributing to virulence based on direct trehalose consumption for increased growth.

Interestingly, we support this claim by observing *ΔscrA/treA* during most assays having an additive effect compared to the single mutants *ΔscrA* or *ΔtreA.* The exception being xylem sap sugar composition modulation assays. Xylem sap previously infected with *ΔscrA* supports more wildtype growth and increases sucrose concentration however xylem sap previously infected with *ΔscrA/treA* supports more wildtype growth than *ΔscrA* parent but fails to increase sucrose concentration as high as *ΔscrA* parent. This is compounded by the finding that *ΔtreA* increase sucrose concentration compared to wildtype. We suggest that small quantities of trehalose are regulating plant carbon sources in host xylem.

Trehalose is known for its roles in plant metabolic regulations and stress tolerance (Lunn et al., 2014, Paul et. al., 2008). Arabidopsis plants use trehalose 6-phoshate as a balancing signal for leaf carbon storage systems, managing sucrose and starch ratios (Kolbe et al. 2005). Among many other metabolites, trehalose is induced 19-fold in infected compare to healthy tomato plants and bacteria treA is upregulated *in planta* during late disease (Lowe-Power et. al. 2016, Jacobs *et al*., 2012). TreA may breakdown trehalose, signaling molecule, to maintain the proper balance required of enrichment of tomato xylem sap without sparking plant defense.

### Link bacterial catabolism, life stage, and virulence

Ralstonia catabolic profile during given infection stages contributes to successful virulence. *Rs* faces niches with diverse and scares nutrient availabilities and used specific carbon pathways to be success if each environment. *Rs* myo-inositol is induce in tomato root exudate and *Rs* used it to gain advantage early in infection (Kravchenko et. al., 2003). During late disease *Rs* utilizes sucrose and trehalose for food and possibility to balance plant responses in the favor of the pathogen (Kolbe et al. 2005). This economically important pathogen strategically takes advantage of myo-inositol, sucrose, and trehalose, compounds the many microbes can consume, and could modulate the environment to be more favorable for *Rs* successful virulence.

## Conclusion

We take a holistic approach to understand the how the host xylem environment shapes bacterial wilt disease caused by *R. solanacearum.* We report evidence that xylem sucrose and trehalose are required for late/stem infection while myo-inositol is requiring for early/root infection. We discovery that *Rs* induces and consumes plant sucrose and that treA may be required for maintaining a constant sucrose concentration inside of host xylem. Together these experiments describe the primary metabolic needs of successful *Rs* cells during early and late infection. This contributes to our understanding of how this devastating plant pathogen is successful in or excluded from the nutritionally harsh xylem and root environment.

## Materials and Methods

### Mutant Construction

#### Standard protocols

Cloning, restriction digestion, sequencing, and PCR were performed using standard methods. R. solanacearum and E. coli were transformed by electroporation or natural transformation as previously described.

#### Services

DNA sequencing were performed at the University of Wisconsin-Madison Biotechnology Center. Oligonucleotide were synthesize by Integrated DNA technologies.

#### Deletions Mutants

Gibson assembly was used to create in-frame deletion using primers listed in Supplemental Table. Deletion constructs were introduced into the chromosome of wild-type *R. solanacearum* strain GMI1000 by double homologous recombination as previously described to create GMI1000ΔiolG, GMI1000ΔscrA, GMI1000ΔtreA, and GMI1000ΔscrA/treA, respectively.

#### Marking unmarked mutants

Natural transformation of plasmids containing Kanamycin or Gentamycin plasmid was used to introduced resistance marker into the chromosome of *R. solanacearum* unmarked mutants. The correct allelic replacement in each mutant and complement was confirmed by PCR and sequencing analyses.

#### Complementation Mutants

Splicing by overlap extension was used whole gene replacement constructs using primers listed in Supplemental Table. Complementation constructs were introduced into the chromosome of respective *R. solanacearum* mutant GMI1000 by double homologous recombination as previously described to create GMI1000ΔiolG.COMP, GMI1000ΔscrA.COMP, GMI1000ΔtreA.COMP, and GMI1000ΔscrA/treA.COMP, respectively.

### Bacterial Growth

#### Media

*R. solanacearum* GMI1000 (WT) was grown in Boucher-Minimal medium broth (BMM), Casamino Acids-Peptone-Glucose rich (CPG) medium liquid broth and plates.

#### Conditions

Plates were grown stagnate at 28°C for 48 hours prior to use. Liquid broth cultures were grown shaking at 28°C.

#### Antibiotics Additives

Antibiotics were added to plates and broth as needed when a marked mutant was being cultured – kanamycin at 0.5 μL/mL of media, spectinomycin at 1 μL/mL of media, gentamicin at 1 μL/mL of media, tetrazolium chloride (TZC) at 2 μL/mL of media. (Modified from Denny & Hayward et al. 2001)

### Catabolic Mutant Media Growth

#### Characterization

##### Growth solutions were prepared

Boucher-Minimal medium broth lacking a carbon source was use as the base solution. 10mM of carbon source (glucose, myo-inositol, sucrose, or trehalose) was add to create sole carbon source growth media.

##### Strains were prepared

Each strain was grown in 3 mL of a diluted 10 mM solution of each media type. The tubes shook in a 28°C incubator overnight to test for the presence or absence of growth on each sole carbon source over 4 biological replicates.

### Catabolic Mutant Xylem Sap Growth

#### Characterization

##### Growth solutions were prepared

Ex vivo xylem sap was harvest as previously describe^5^. However, these plants were pre-inoculated with GMI1000 Wildtype, ΔiolG, ΔscrA, ΔtreA, or ΔscrA/treA which creates five types of ex vivo xylem sap that experienced different *R. solanacearum* catabolic potentials. Ex vivo sap used was normalized to bacterial colonization of 1×10^5^ CFU/gram.

##### Strains growth

A Titertek Multiskan spectrophotometer plate reader measuring AB600 was used to quantify bacterial growth in each media type for 24 hours. Bacterial suspensions of each strain were added to 50ul of each type of *ex vivo* xylem sap growth media in 96-well, flat-bottom, half-area plates. Bacterial suspensions were adjusted to an OD of 0.01. Absorbance curves where produce and Mann-Whitney analysis were done of area under curve values for each strain compared to wildtype.

### Plant Virulence Assay

#### Plants growth

Wilt-susceptible tomato plants (*Solanum lycopersicum*) cultivar Bonny Best (BB) were used for *in planta* virulence assays. BB were sown and watered every day for 14 days to promote healthy germination. The 14-day old plants were then transplanted and watered with Hoagland’s Solution and water in an alternating pattern until one day prior to inoculation.

#### Soil soak bacterial inoculation

On the 17th day, an unwounded plant were inoculated using a naturalistic soil soak protocol. Each plant was soaked with a 50 mL inoculum of bacterial suspension, adjusted to 10^7 CFU/mL. As a negative control, 15 BB plants of each biological replicate were inoculated with 50 mL of DI water.

#### Rating Disease

The disease progress of each plant was rated daily for 14 days post-inoculation (dpi) on a 0-4 disease index scale – where 0 represented no symptoms, 1 corresponded to 1-25% leaf area wilting, 2 for 26-49%, 3 for 50-74%, and 4 for 75-100% of the leaf area wilted. These virulence assays were repeated for 4 independent biological replicates and were analyzed through repeated measures of ANOVA.

### *In Planta* Colonization & Competition

#### Stem Inoculations

Plants for colonization are inoculated with either the wild-type strain or the mutant using the soil soak method as described above. Plants for competition are inoculated with marked wildtype and marked mutant cell suspensions were combined in a 1:1 ratio using the petiole method as described. On the 21^st^ day, cut petiole plant were inoculated. Each plant was inoculated with a 2000 cell Bacterial suspensions. As a negative control, 15 BB plants of each biological replicate were cut and inoculated with DI water.

#### Root Inoculations

Plants were inoculated with either the wild-type strain or the mutant using the root attachment method as previously described^7^.

Plants were inoculated with marked wildtype and marked mutant cell suspensions were combined in a 1:1 ratio using the root attachment method as described above.

#### Sterilization

Stem colonization and competition require no extra surface sterilization. Roots are sterilized by 10% bleach for 30 seconds, 70% ethanol for 30 seconds, and sterilize water for 1mins.

#### Quantification

Stem population sizes of each strain in tomato plants were quantified as described. Briefly, at a given time points after inoculation, plants inoculated with each strain were segmented ∼1cm, weighed, and ground in 1 ml sterile deionized water. Root population sizes of strain in tomato plants were quantified by collecting root at different time points after inoculation. Two whole seedling roots are weighed and ground in 300 ul sterile deionized water. The resulting homogenate was dilution plated on TZC plates supplemented with 100 mg l-1 cycloheximide and appropriate antibiotics. R. solanacearum population sizes were determined as cfu g-1 of plant tissue. Methods modified from Khokhani et. al., 2018. All experiments were repeated at least three times. Competitive Index was calculated and analysis base on previously described methods^8^. Statistical Analysis is done using Mann–Whitney U test or Wilcoxon sign-rank test.

#### *Ex vivo* xylem sap collection

Xylem sap was harvested from plants 5 days post inoculation after. Plants to ensure adequate water status, plants were well watered each evening before sampling. Plants inoculated by soil soaking were detopped 2cm above the cotyledons, and plants inoculated by petiole inoculation were detopped at the site of inoculation. Root pressure allowed xylem sap to accumulate on the stump. The sap accumulated in the first 2-3 min was discarded and the stump was washed with water and gently blotted dry with a Kimwipe. Over 30 min, the accumulating sap was frequently transferred into pre-chilled 1.5 ml flip cap microcentrifuge tubes held in -20°C block or on ice. Samples were flash-frozen and stored at -80°Cuntil analysis. Subsequently,100 mg stem samples were harvested and stem colonization was quantified as stated above. (Modified from Lowe-Power et. al. 2018)

#### GOD/invertase enzymatic assay to determine sucrose concentration

in a 96-well plate, 85 ul distilled water, 5 uL from each sample and 10 uL invertase were placed in each well. The invertase was prepared at a concentration of 10 mg/mL in distilled water. The plate was then sealed and placed in a water bath at 55 C for 10 min. After this period the plate was removed from the water bath and 200 lL of GOD (Bioclin kit) reagent was added, and the plate was sealed again and placed in a water bath at 37 C for 15 min. After this time, the plate was removed from the water bath and placed at room temperature for 5 min. The absorbance at 490 nm was read in a Titertek Multiskan Plus spectrophotometer, equipped for reading plates. Standard sucrose solutions were also added to each plate in separate wells at concentrations 10-1000 umol sucrose. This allowed a calibration curve to be constructed which was used to determine the sucrose concentration in each sample. Each analysis was done in triplicate. (Modified from Teixeira et. al. 2012)

## Funding

This research was supported by the University of Wisconsin-Madison College of Agricultural and Life Sciences, a UW-Madison Sophomore Research Award to OS and a SciMed-GRS Fellowship to CGH.

## Acknowledgements

The authors thank Devanshi Khokhani and Tiffany Lowe-Power for helpful discussions.

## References

Ailloud, F., Lowe, T., Cellier, G., Roche, D., Allen, C., and Prior, P. (2015). Comparative genomic analysis of Ralstonia solanacearum reveals candidate genes for host specificity. BMC Genomics 16:270. doi: 10.1186/s12864-015-1474-8

Allen, C., Prior, P., and Hayward, A.C., (2005). Bacterial Wilt Disease and the Ralstonia solanacearum Species Complex (St. Paul, MN: APS Press).

Álvarez, B., E. G. Biosca, and M. M. López. 2010. On the life of Ralstonia solanacearum, a destructive bacterial plant pathogen, p. 267–279. In A. Mendez-Vilas (ed.), Current research, technology and education topics in applied microbiology and microbial biotechnology, vol. 1. Formatex, Badajoz, Spain

Bais HP., Weir TL., Perry LG., Gilroy S., Vivanco JM. (2006) The role of root exudates in rhizosphere interactions with plants and other organisms. Annu Rev Plant Biol 57: 234–266.

Caldwell, D., Kim, B. S., and Iyer-Pascuzzi, A. S. (2017). *Ralstonia solanacearum* differentially colonizes roots of resistant and susceptible tomato plants. Phytopathology 107, 528–536. doi: 10.1094/PHYTO-09-16-0353-R

Cepulyte R, Danquah WB, Bruening G, Williamson VM (2018) Potent attractant for root-knot nematodes in exudates from seedling root tips of two host species. Sci Rep 8(1):10847

Chellemi, D., Andersen, P., Brodbeck, B., Dankers, W., and Rhoads, F. (1998) Correlation of chemical profiles of xylem fluid of tomato to resistance to bacterial wilt. In Bacterial Wilt Disease. Berlin, Germany: Springer, pp. 225–232.

Coplin DL, Sequeira L, Hanson SR. 1974. Pseudomonas solanacearum: virulence of biochemical mutants. Canadian Journal of Microbiology 20:519–529.

Dekel E, Alon U (2005) Optimality and evolutionary tuning of the expression level of a protein. Nature 436: 588–592

Denny T, Hayward AC. 2001. Ralstonia. In Laboratory Guide for Identification of Plant Pathogenic Bacteria, 3rd Ed., ed. N Schaad, JB Jones, W Chun, pp. 165–89. St. Paul, MN: APS Press

Dixon, G.R. and Pegg, G.F. (1972) Changes in amino-acid content of tomato xylem sap following infection with strains of Verticillium albo-atrum. Ann. Bot. 36, 147–154

Elphinstone, J. G. 2005. The current bacterial wilt situation: a global overview, p. 9–28. In C. Allen, P. Prior, and A. C. Hayward (ed.), Bacterial wilt disease and the Ralstonia solanacearum species complex. APS Press, St. Paul, MN.

Evert, R. and Eichhorn, S. (2006) Esau’s Plant Anatomy: Meristems, Cells, and Tissues of the Plant Body: Their Structure, Function, and Development, Wiley

Fatima, U. and Senthil-Kumar, M. (2015) Plant and pathogen nutrient acquisition strategies. Front. Plant Sci. 6, 750 27.

Genin, S. and Denny, T.P. (2012) Pathogenomics of the Ralstonia solanacearum species complex. Annu. Rev. Phytopathol. 50, 1–23

Gnanamanickam, S. S. (2006) Plant-Associated Bacteria 1–712, Printed in the Netherlands

Gudelj I, Beardmore RE, Arkin SS, MacLean RC (2007) Constraints on microbial metabolism drive evolutionary diversification in homogeneous environments. J Evol Biol 20: 1882–1889

Hayward, A. C. 1991. Biology and epidemiology of bacterial wilt caused by Pseudomonas solanacearum. Annu. Rev. Phytopathol. 29:67–87.

Jacobs, J. M., Babujee, L., Meng, F., Milling, A., and Allen, C. (2012) The in planta transcriptome of Ralstonia solanacearum: conserved physiological and virulence strategies during bacterial wilt of tomato. mBio 3, e00114–12

Khokhani, D., Lowe-Power, T., Tran, T., Allen, C. (2017) A single regulatory switch in a plant pathogen mediates strategic trade-offs between growth or virulence and attachment or spread. mBio 8, e00895–17

Khokhani, D., Tran, T., Lowe-Power, T., Allen, C. (2018) Plant Assays for Quantifying Ralstonia solanacearum Virulence. Bioprotocols 8: 18

Kolbe, Anna, Axel Tiessen, Henriette Schluepmann, Matthew Paul, Silke Ulrich, and Peter Geigenberger. 2005. “Trehalose 6-Phosphate Regulates Starch Synthesis via Posttranslational Redox Activation of ADP-Glucose Pyrophosphorylase.” Proceedings of the National Academy of Sciences of the United States of America 102 (31): 11118–23. https://doi.org/10.1073/pnas.0503410102.

Kravchenko LV, Azarova TS, Leonova-Erko EI, Shaposhnikov AI, Makarova NM, Tichonovich IA (2003) Root exudates of tomato plants and their effect on the growth and antifungal activity of *Pseudomonas* strains. Microbiology 72:37–41

Lowe-Power TM, Khokhani D and Allen C, 2018. How Ralstonia solanacearum exploits and thrives in the flowing plant xylem environment. Trends in Microbiology, 26, 929–942.

Lowe-Power, T.M., Hendrich, C.G., von Roepenack-Lahaye, E., Li, B., Wu, D., Mitra, R., Dalsing. B, Ricca P, Naidoo J, Cook D., Jancewicz A., Masson P,. Thomma B., Lahaye T., Michael A., Allen C., (2018) Metabolomics of tomato xylem sap during bacterial wilt reveals Ralstonia solanacearum produces abundant putrescine, a metabolite that accelerates wilt disease. Environ. Microbiol. 20, 1330–1349

Lowe-Power, T., Khokhani, D., Allen, C. (2018b) How Ralstonia solanacearum Exploits and Thrives in the Flowing Plant Xylem Environment. Trends in Microbiology. 26(11): 929–942

Lunn, John Edward, Ines Delorge, Carlos María Figueroa, Patrick Van Dijck, and Mark Stitt. 2014. “Trehalose Metabolism in Plants.” Plant Journal 79: 544–67. https://doi.org/10.1111/tpj.12509.

McGarvey J, Denny TP, Schell MA. 1999. Spatialtemporal and quantitative analysis of growth and EPS production by *Ralstonia solanacearum* in resistant and susceptible tomato cultivars. Phytopathology 89:1233–39

Paul, Matthew J, Lucia F. Primavesi, Deveraj Jhurreea, and Yuhua Zhang. 2008. “Trehalose Metabolism and Signaling.” Annual Review of Plant Pathology, no. 59: 417–41. Plant Physiol. 99: 1267–70

Teixeira et al., (2012) Development of a method to quantify sucrose in soybean grains. Food Chem., 130 (2012), pp. 1134–1136

Thines E, Weber RW, Talbot NJ (2000) MAP kinase and protein kinase A-dependent mobilization of triacylglycerol and glycogen during appressorium turgor generation by Magnaporthe grisea.. Plant Cell. 2000 Sep; 12(9):1703–18.

Wang, E.L. and Bergeson, G.B. (1974) Biochemical changes in root exudate and xylem sap of https://doi.org/10.1146/annurev.arplant.59.032607.092945.

Paul, Matthew J, Lucia F. Primavesi, Deveraj Jhurreea, and Yuhua Zhang. 2008. “Trehalose Metabolism and Signaling.” Annual Review of Plant Pathology, no. 59: 417–41. https://doi.org/10.1146/annurev.arplant.59.032607.092945.

Pfeiffer T, Bonhoeffer S (2004) Evolution of crossfeeding in microbial populations. Am Nat 163: E126–E135

Pollock CJ, Farrar J. 1996. Source-sink relations: the role of sucrose. In Environmental Stress and Photosynthesis, ed. NR Baker. The Netherlands: Kluwer. In press

Polz MF, Cordero OX. 2016. Bacterial evolution: genomics of metabolic trade-offs. Nat Microbiol. 1:16181

Prior, P., Ailloud, F., Dalsing, B. L., Remenant, B., Sanchez, B., and Allen, C. (2016). Genomic and proteomic evidence supporting the division of the plant pathogen Ralstonia solanacearum into three species. BMC Genomics 17:90. doi: 10.1186/s12864-016-2413-z

Salanoubat M, Belliard G. 1989. The steady-state level of potato sucrose synthase mRNA is dependent on wounding, anaerobiosis and sucrose concentration. Gene 84: 181–85

Siebrecht S, Herdel K, Schurr U, Tischner R. Nutrient translocation in the xylem of poplar: diurnal variations and spatial distribution along the shoot axis, Planta, 2003, vol. 217 (pg. 783-793)

Sonnewald W, Willmitzer L. 1992. Molecular approaches to sink-source interactions. tomato plants infected with Meloidogyne incognita. J. Nematol. 6, 194–202 26.

White, M. C., Chaney, R. L. and Decker, A. M. 1981. Metal complexation in xylem fluid. III. Electrophoretic evidence. Plant Physiol., 67: 311–315.

Yang CH, Crowley DE. (2000) Rhizosphere microbial community structure in relation to root location and plant iron nutritional status. Appl Environ Microbiol. 66:345–351. doi: 10.1128/AEM.66.1.345-351.2000.

Zuluaga, A. P., Puigvert, M., and Valls, M. (2013). Novel plant inputs influencing *Ralstonia solanacearum* during infection. Front. Microbiol. 4:349. doi: 10.3389/fmicb.2013.00349

